# CARDIAX-NNFE A Cardiac Mechanics SciML Framework

**DOI:** 10.64898/2026.07.27.741090

**Authors:** Benjamin J. Thomas, Michael S. Sacks

## Abstract

One goal of Scientific Machine Learning (SciML) is to advance traditional scientific computing frameworks with modern machine learning tools. This includes extending established methods, such as the finite element method, with cardiac function applications due to their complexity and need for very rapid execution times for real time clinical use. In this work, we present an advanced form of the Neural Network Finite Element (NNFE) method specialized for cardiac simulations, termed `CARDIAX-NNFE`. The NNFE method learns the parameter-to-displacement field map by training over the residual of the hyperelastic material PDE, using the domain represented by finite elements. The implementation is developed in Python using `JAX` to leverage its automatic differentiation, highly parallel GPU, and JIT-compilation capabilities. To demonstrate `CARDIAX-NNFE` effectiveness, we trained full cardiac pressure-volume responses using a simplified heart model, spanning the entire cardiac physiological functional range. Results indicated the ability to simulate a family of pressure-volume solutions with average nodal positional error of 0.023 mm and maximal error of 0.054 mm, with a single complete PV loop evaluated in 0.002 seconds. The CARDIAX-NNFE software platform thus provides for a robust platform for cardiac functional simulations. Moreover, it provides the structure for residual-based SciML methods, which can apply to a variety of physics-based biomedical problems that require high execution speed for clinical applications.

## Metadata

**Table 1:** Code metadata.

| Nr. | Code metadata description | Metadata |
| --- | --- | --- |
| C1 | Current code version | 0.1.0 |
| C2 | Permanent link to code/repository used for this code version | https://github.com/WCCMS-UTAustin/CARDIAX-NNFE |
| C3 | Permanent link to Reproducible Capsule | none |
| C4 | Legal Code License | GNU General Public License (GPL) |
| C5 | Code versioning system used | git |
| C6 | Software code languages, tools, and services used | Python, JAX |
| C7 | Compilation requirements, operating environments & dependencies | Linux, CUDA |
| C8 | If available Link to developer documentation/manual | https://WCCMS-UTAustin.github.io/CARDIAX-NNFE/ |
| C9 | Support email for questions |  |

## 1. Motivation and significance

Research into machine learning for solving PDEs has increased significantly in the past decade, but had started quite some time ago [8, 9]. While machine learning has shown great success in many fields, its general adoption has been slow in many scientific computing fields because of the lack of proper and robust methods development. Scientific Machine Learning (SciML) has thus emerged to bridge the gap between machine learning and traditional scientific computing. The goal of SciML is to leverage the strengths of machine learning, such as its approximation power and fast evaluations, while maintaining the reliability and accuracy of traditional scientific computing methods. In addition, for well established methods such as the finite element (FE), a more modern approach is to utilize a *differentiable* framework [16, 13]. However, a differentiable code is only half the battle, as while it provides the foundation for a SciML approach, it leaves all machine learning integration tasks to the user.

A key area where SciML can be quite impactful is computational medicine, wherein the governing equations are very complex, nonlinear, multiscale, and multiphysics, but extreme computational speed is also essential for clinical use. A prime example is modeling heart in the normal and diseased states (e.g. myocardial infarction). Accurate cardiac computational simulations can play a pivotal role in improving our understanding of the cascade of events that lead to progressive heart failure, as well as guide future therapies. However, computational speed remains a major bottleneck preventing routine clinical translation, as well as more physiologically realistic models. Addressing these challenges can open the door for ‘digital twins’, a modeling approach based on continuously updating patient-specific data to better predict the future state of the patient [6].

Herein, we will demonstrate a codebase that is both high performance and also merges a differentiable FE framework with modern machine learning tools, termed the Neural Network Finite Element (NNFE) method. The initial description of the NNFE method was reported in Zhang et. al [10] to learn the boundary condition-to-displacement map describing the mechanics of myocardium tissue through an energy minimization approach. This method was latter utilized for heart valve simulations that included leaflet contact [11]. To handle more complex problems, the method was reframed to be residual-based to handle cardiac simulations over the entire physiological range [12], unlocking the cardiac digital twin potential. The NNFE method now learns the parameter-to-displacement map by computing the PDE residual with the FE discretization [16]. Critically, the residual is treated as the loss function so the error metric is the same as traditional FE methods. Moreover, the data-independent training is very helpful in data scarce areas such as clinical applications where many-query results are desired. In this work, we demonstrate a codebase that allows for the required flexibility to extend to more realistic cardiac models while maintaining the ease of use needed to make SciML more accessible to the broader academic community.

## 2. Software description

The best design choice for the software is to use `JAX` because of its vectorization, automatic differentiation, and JIT-compilation capabilities. Utilizing `JAX` we built the FE framework using `CARDIAX` along with `Equinox` and `Optax` for the machine learning components. The software handles the orchestration of these packages to create the necessary functions to train a network in a fully jittable function, managing the FE state while allowing the ML functionality. A more in depth mathematical description of the method to complement the orchestration is given in Appendix A. The abstractness of the method is the key to its flexibility to handle more complex problems.

### 2.1. Software architecture

The software is structured in a modular manner that allows for flexibility across the vast amount of components that can be modified when combining finite elements with machine learning. The main object of interest is the `NNFEManager` which is responsible for combining the various components appropriately and is the only object that the user needs to interact with. This object is created through combining a number of smaller Managers which can be run independently. Thus, each individual manager handles its own state and can be modified as desired without affecting the overall structure of the code.

The first block is the yaml file parsing which highlights the reproducibility of the framework. A master yaml file can be passed as input to the from config method of the `NNFEManager` which will parse the files into frozen dataclasses representing the configuration files for each manager. The initializations can either go through the standard yaml file parsing or can be done through a dictionary if the user wants to customize further than what the standard functionality allows for. Most assertions are done at the parsing stage which allows for errors to be thrown quickly before any utility is actually being done. The frozen dataclass then prevents any accidental modifications that may be made which can throw errors or cause reproduciblity issues.

After configurations are parsed appropriately, the NNFE Factory ensures that each config creates the appropriate manager. The factory creates four main objects:

- FEManager: Creates and holds the state of the PDE for assembly
- MLManager: Create the neural networks and optimizer used for training.
- ProjectManager: Prints training stats, creates directories, checkpoints, and saves results.
- Sampler: Samples the training and testing space.

These managers have dependencies on each other that are handled in the factory like DoFs from the FEManager needs to be the output size of the neural network in the MLManager. These interdependencies are set in the yaml config files to again ensure reproducibility.

The crux of this framework is the `NNFEManager` which pulls the above managers together and creates the methods to be used by the users. One of the most important components is the look at the `NNFEParamsConfig` which defines the parameters the network will train over, marking them as functional as opposed to static. This happens for both essential and natural boundary conditions even though these are treated in very different ways when the residual is computed. This residual is then differentiated with respect to the network parameters to compute the gradient used for optimization, and a `make_step` method is used for training. A predefined `train` loop is created to run the optimization procedure, but the user can create their own if preferred. Along with the construction of the training process, a corresponding `test` procedure is created to evaluate the trained model on a testing set. Lastly, `config dump` is used to save the master yaml file if used, and a `save` method for saving the model and optimizer states.

### 2.2. Software functionalities

Most functionalities of the software are derived from the functionalities of the various packages used. In this manner, we obtain the most flexibility by inheriting all FE capabilities from `CARDIAX` and all the machine learning capabilities from `Equinox` and `Optax`. The abstraction of the yaml files allow for this high flexibility at the manager level. This means if a PDE can be solved with FE in `CARDIAX`, then the functionality exists to attempt to train a network to solve it. The same goes for the numerous machine learning architectures that can be created in `Equinox` and the various optimizers allowed in `Optax`.

The four main functionalities that the user will interact with beyond the configuration files are:

- `train`: Runs the main training loop while saving stats to track progress and times itself to compare runtimes. A description of the construction is shown in Figure 2.
- `test`: Evaluates the trained model on a testing set by generating traditional FE solutions where the network prediction is used as the initial guess to the solver
- `save`: Saves the current state of the model and optimizer to a checkpoint file for continued training and evaluation
- `config dump`: Saves the master yaml file that was used to create the nnfe object

**Figure 1:**
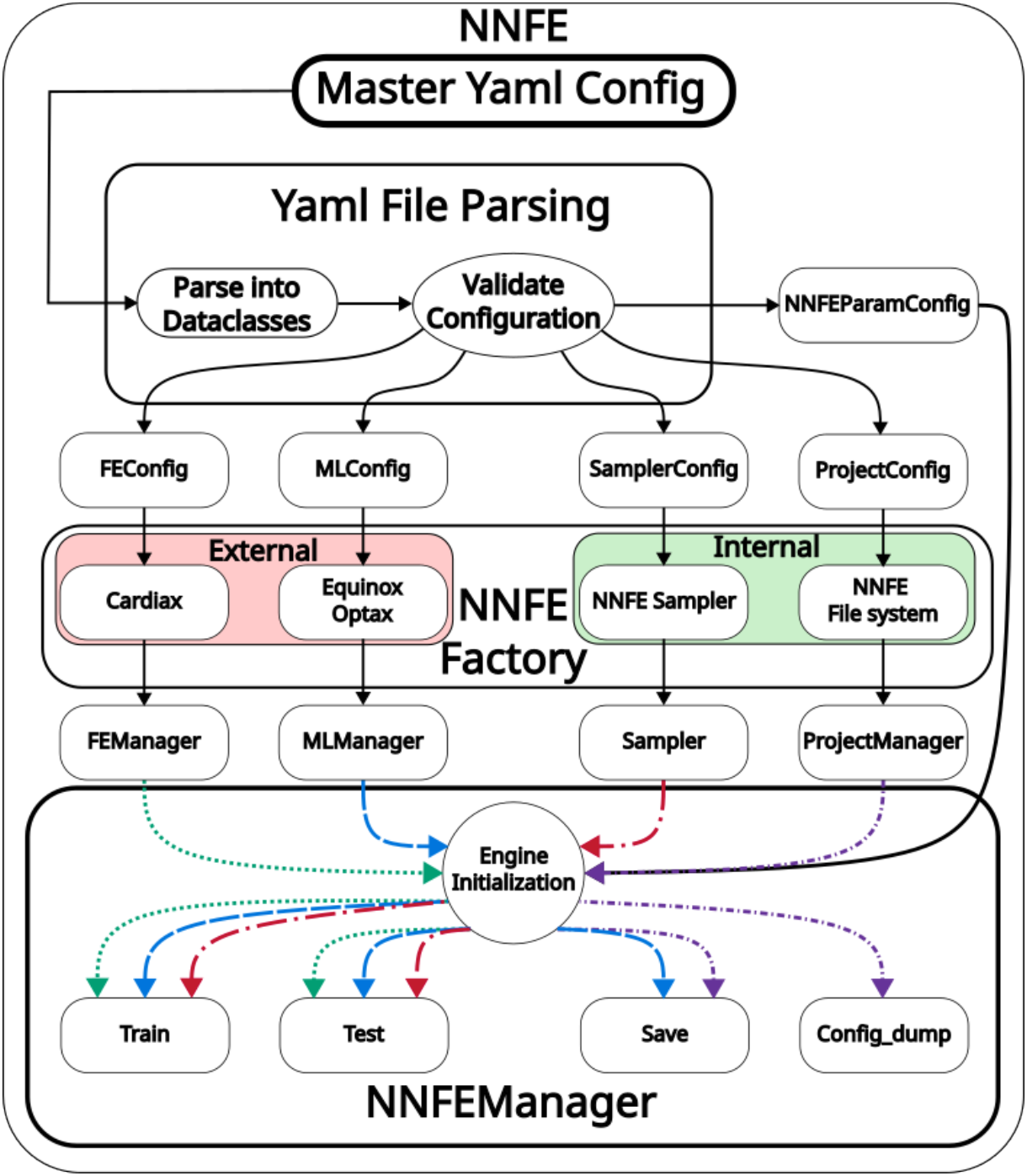
The flowchart shows the architecture of the codebase. The user provides the Master Yaml Config that is parsed into various configuration dataclasses. These data-classes go into the NNFE Factory responsible for spawning the various managers. The NNFE Managers then pool together to create the NNFEManager where the colored arrows are used to show the dependency injections.

**Figure 2:**
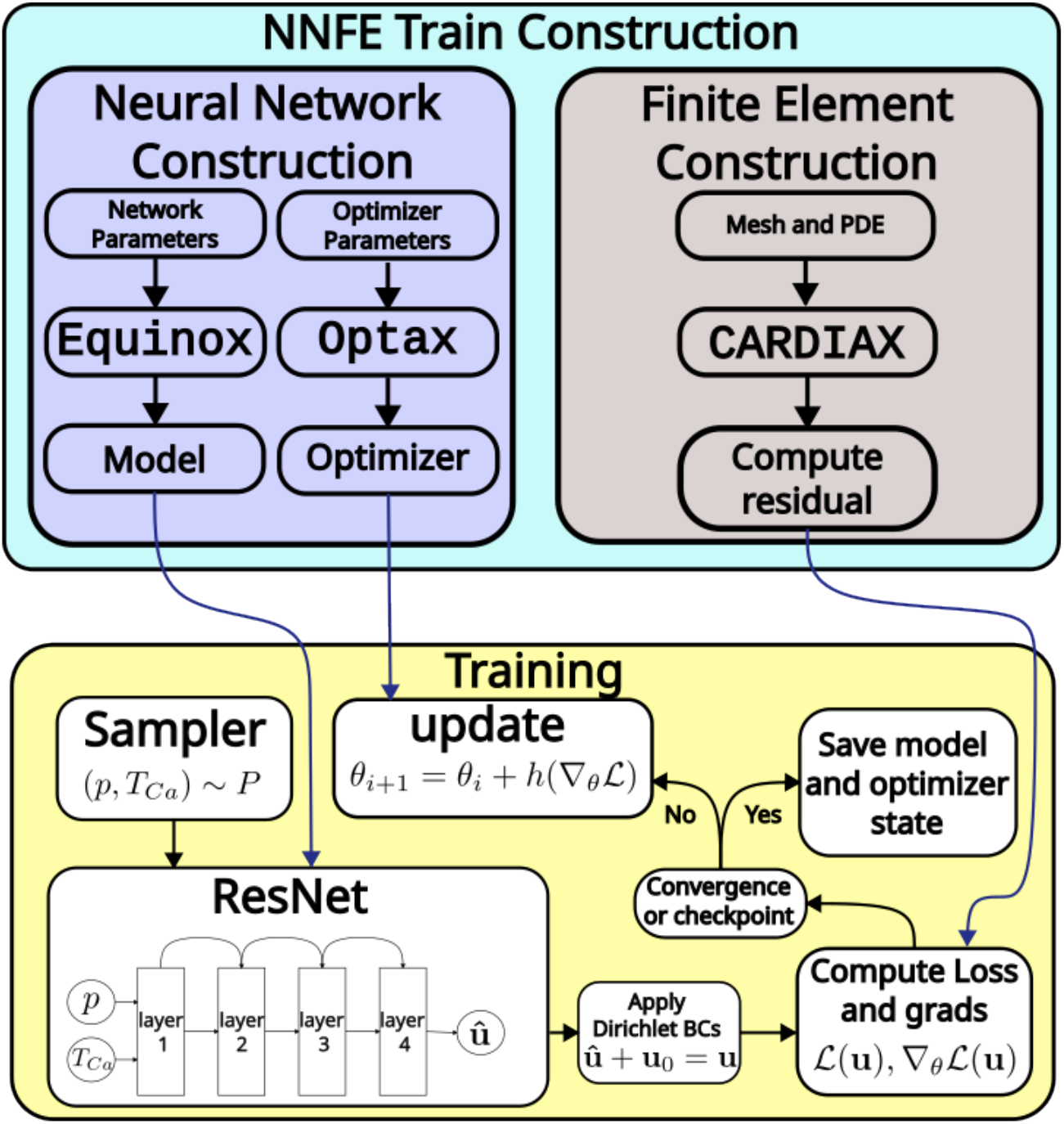
A diagram displaying the construction and content of the `train` method. The network is constructed using `Equinox`, the optimizer with `Optax`, and the residual through `CARDIAX`. These are then combined to form the training loop where samples are drawn, DoFs are predicted, the loss and gradients are computed, and then parameters are updated.

## 3. Illustrative examples

To demonstrate how the code works, we will look into a simplified representation of the left ventricle of the heart, a prolate spheroid (PS). The cardiac mechanics for the PS are nonlinear hyperelastic with anisotropy defined through fiber directions [15]. The inner surface is pressure loaded, and an active stress component is used to simulate the contractile forces along the fiber directions. The goal is to learn displacements given the pressure and active stress. We will setup, train, and evaluate the NNFE method for this problem to demonstrate the capabilities of the software. For a more detailed description of the PDE, mesh, and fibers, see Appendix B.

### 3.1. Defining the Finite Element Problem

Given an appropriate mesh, we need to define the FE config file. First, we define the finite element field, *u*, to solve over the PS Fig 3A. After the finite element field is defined, we must select the appropriate PDE [14]. As shown in Figure 3B, the fibers are defined through the mesh while active stress and pressure are defined as constants. The PS 3C is fixed in all directions at the base.

**Figure 3:**
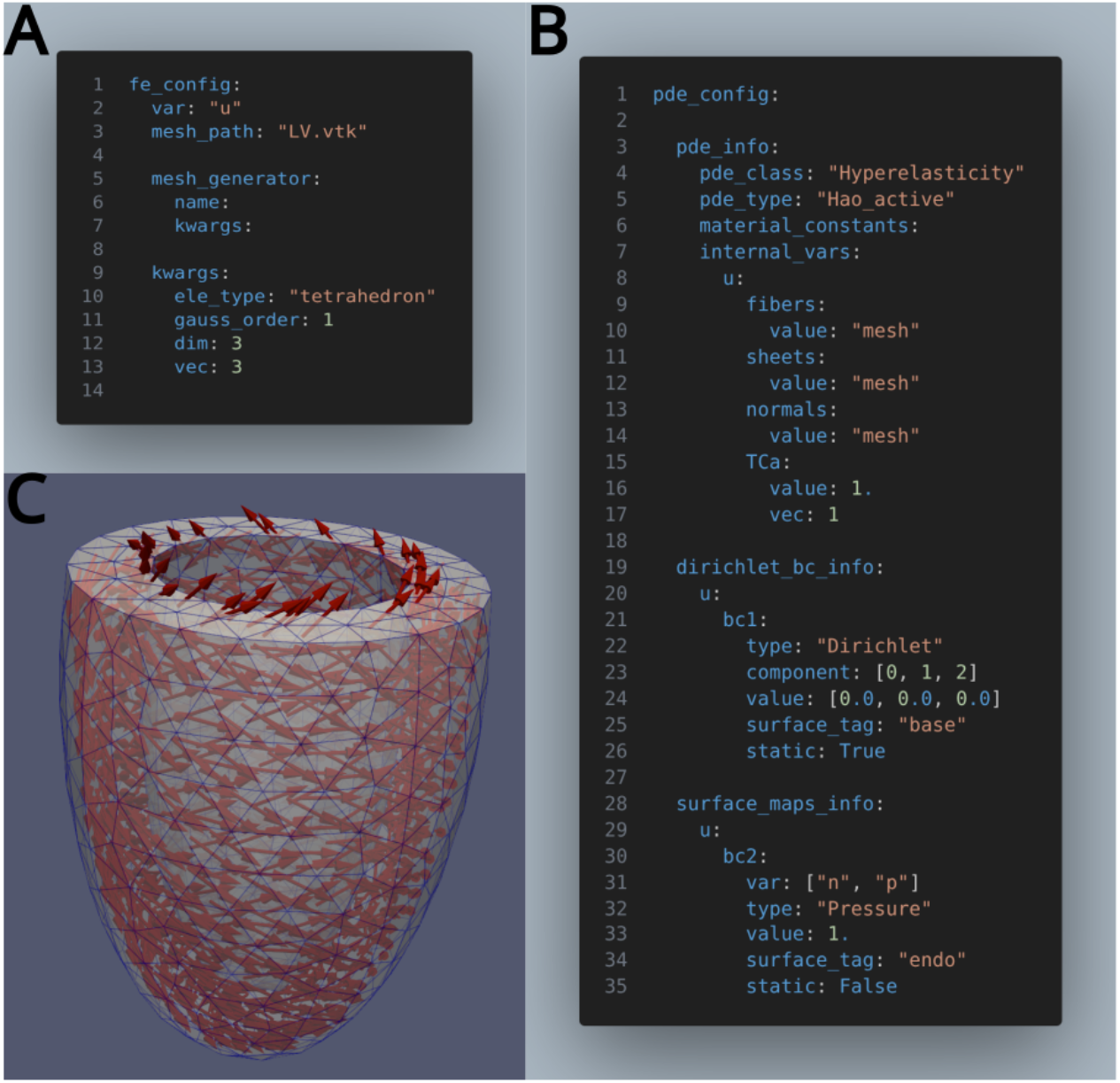
A visualization of the prolate spheroid problem. The geometry is a simplified representation of the left ventricle of the heart. The inner surface is pressure loaded, and an active stress component is used to simulate the contractile forces along the fiber directions. The goal is to learn displacements given the pressure and active stress.

### 3.2. Creating the Neural Network

To create the neural network, we specify the network architecture by the name then kwargs are unpacked to create the desired instance Figure 4A. In this example, we created a ResNet using 6 layers of 128 wide. Importantly, the output size of the network must match the degrees of freedom of the finite element field. The optimizer is created in a similar way where the name of the optimizer is given along with the kwargs to create the desired instance as well as the necessary kwargs for a learning rate scheduler 4B.

**Figure 4:**
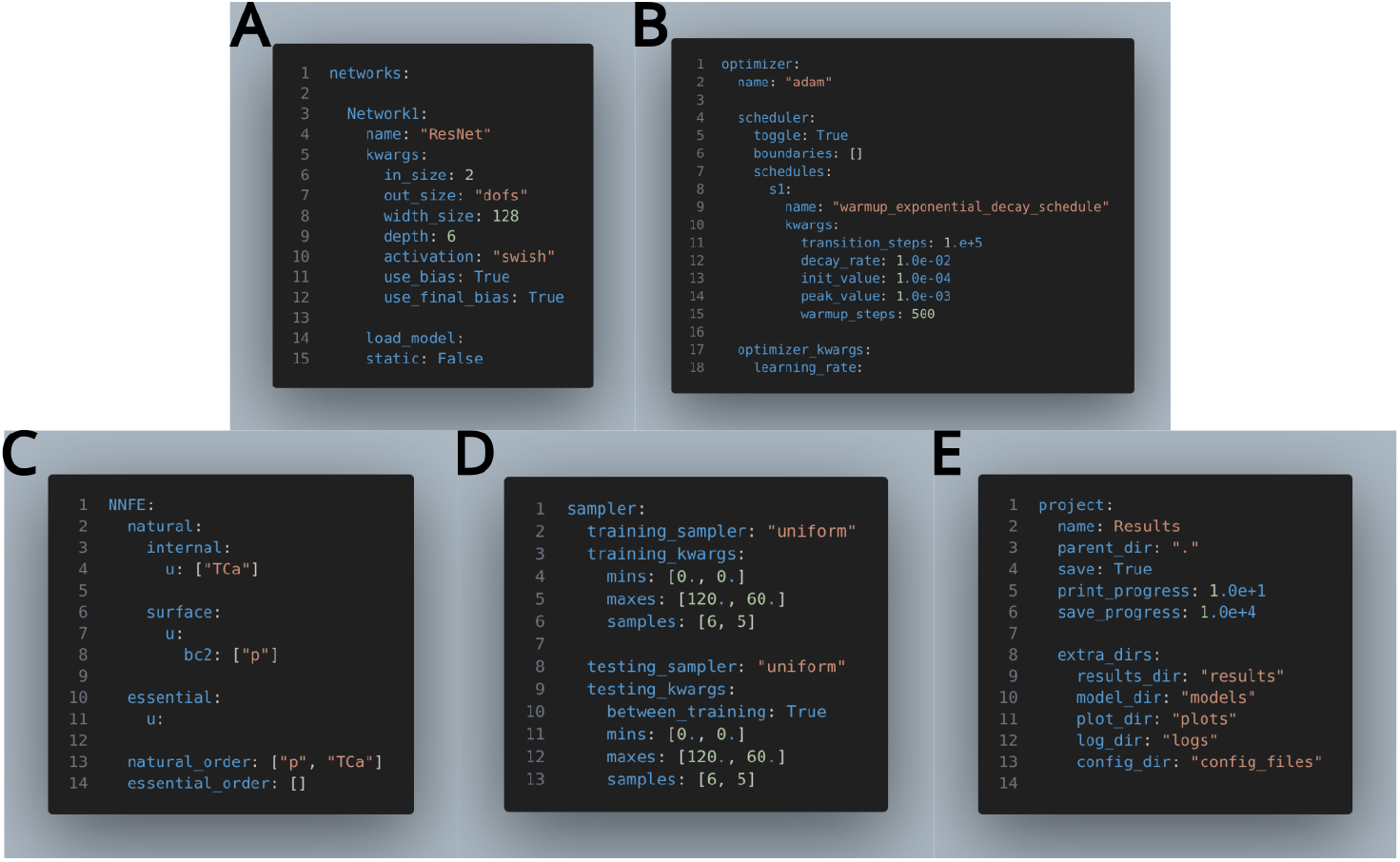
A visualization of the prolate spheroid problem. The geometry is a simplified representation of the left ventricle of the heart. The inner surface is pressure loaded, and an active stress component is used to simulate the contractile forces along the fiber directions. The goal is to learn displacements given the pressure and active stress.

### 3.3. The NNFE Method

The final step is to designate the physiological parameters of the PDE. In this case, we have the external pressure and the magnitude of active stress 4C, so we can evaluate PV-loops in milliseconds after training instead of using traditional FE methods. Since we are choosing two constants fields to train, we pick to uniformly sample from a grid describing the physiological range of each parameter 4D. We also specify where we would like to save checkpoints and final results along with the config files used to create the training procedure, emphasizing reproducibility 4E.

### 3.4. Results

After we `train`, we would like to review the results. We already have the value of the residual from the computed loss. The power of the framework is we can now generate traditional FE solutions through the same configuration file used in the training process and perform FE simulations. Given the speed of the software, we can create a grid of testing points Figure 5 to evaluate pointwise residuals and errors to plot surfaces. We then evaluate the PV loop which lives on this grid. We have greatly improved upon the original version of the codebase in terms of training time, accuracy, and evaluation time Figure 6.

**Figure 5:**
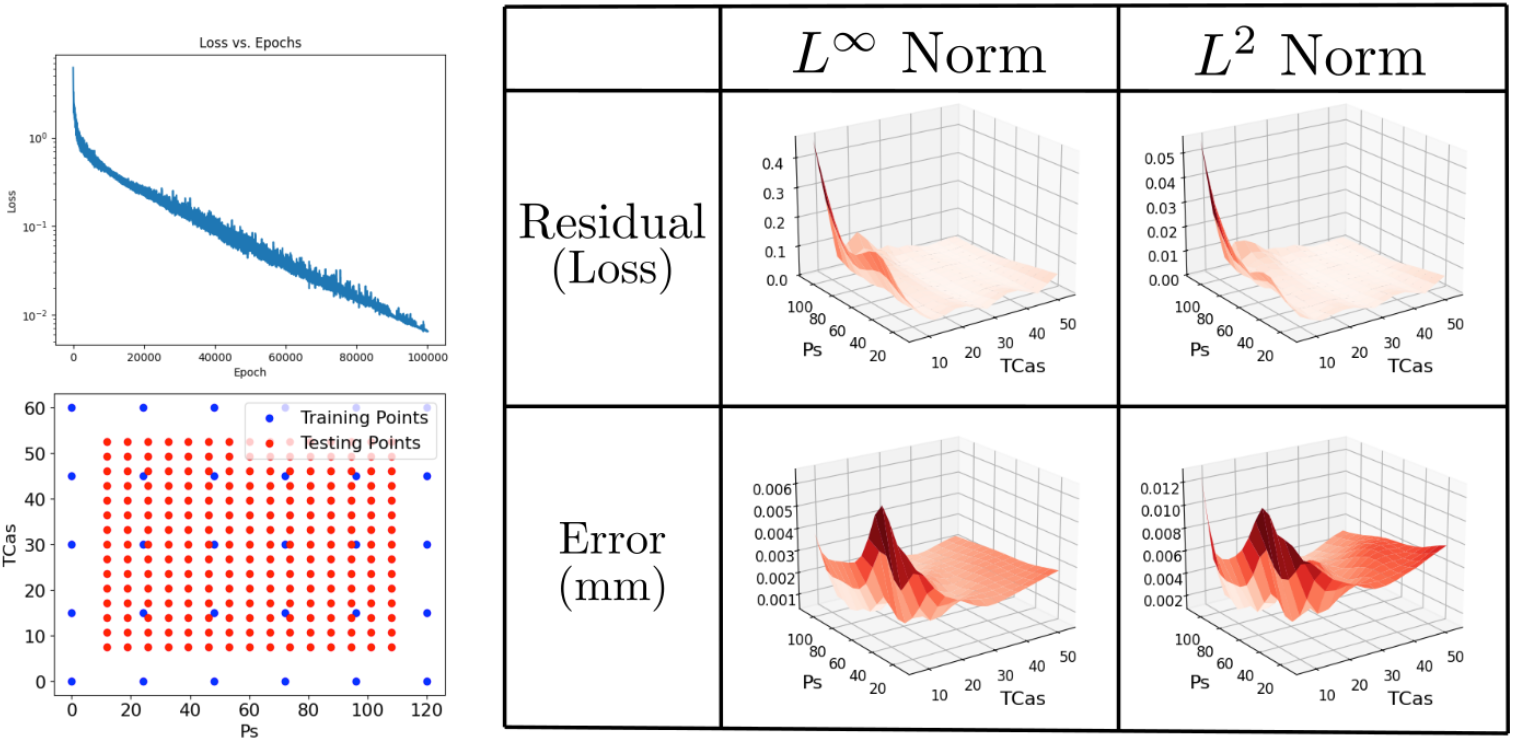
The graph at the top left is the training curve that is the mean of residual losses. The bottom left plot shows the training points in blue to train the network and testing points in red to evaluate the performance on the right. The right plot shows the *L*^2^ and*L*^∞^ residuals and errors vs. FE solutions for the testing points.

**Figure 6:**
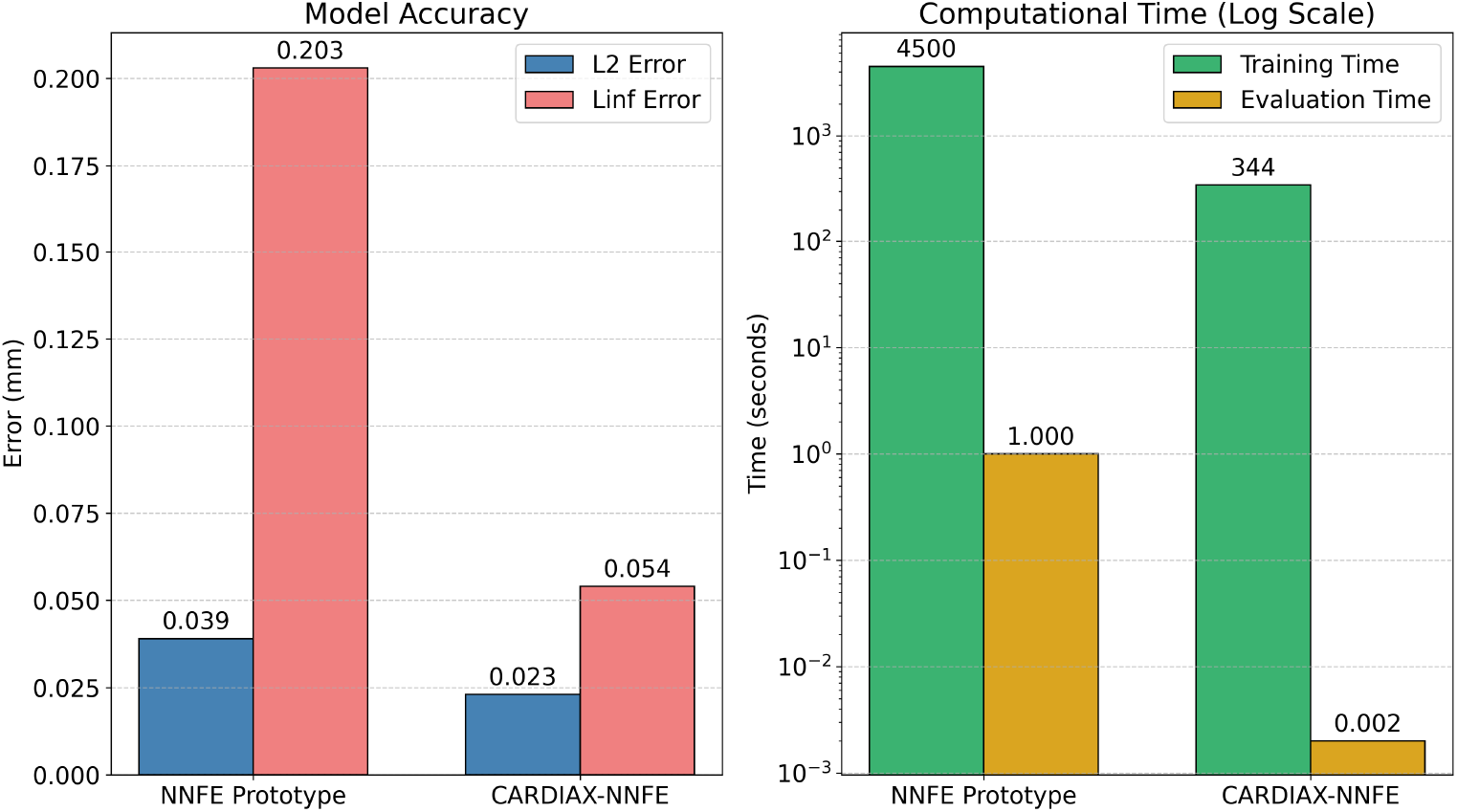
A comparison between the previous version of the code used in [12] and the current code. Average (*L*^2^) and maximal (*L*^∞^) errors are shown on the left with training/evaluation times on the right.

## 4. Impact

The major impact of this software is to provide a framework for NNFE, which is a specific scientific machine learning method. While we demonstrated on cardiac mechanics, the software can be used on a range of problems that can be solved with minimum residual methods. A driver for the acceptance of the software is the ease of use offered by the configuration files. The modular nature of the code allows for custom changes and implementations to be easily added if the user desires. In addition to increasing access to scientific machine learning methods, the software enables the rapid prediction of solutions. A major barrier for clinical research is the ability to obtain results in a clinically relevant timeframe. Many of the traditional finite element methods, while accurate, can take hours to days to compute solutions. By expediting the solution time to the order of seconds or minutes, it would allow for computational models to enter the realm of clinical translational as opposed to purely research tools. This software provides a solid foundation to build upon to achieve the goals required for clinical translation.

## 5. Conclusions

We have created a scientific machine learning software, `CARDIAX-NNFE` that merges minimum residual methods derived from finite elements with machine learning capabilities. We have abstracted the components to a level where the code is modular and flexible to allow for custom implementations. The structure of the code can then be accessed through configuration files, making the code more user friendly and reproducible. We have demonstrated the capabilities of the software by learning a pressure-volume loop of a simplified heart geometry and comparing the final results to traditional finite element solutions. The software provides the foundation needed to further explore scientific machine learning methods and their potential to assisting clinical translation.

## Acknowledgements

*Optional. You can use this section to acknowledge colleagues who dont qualify as a co-author but helped you in some way*.

## Appendix A Mathematical Foundation of the NNFE Method

To utilize the NNFE method, a neural network is first created to map boundary conditions and internal variables of the PDE to the solution field. Since we are focused on cardiac mechanics, we focus on boundary conditions, as the model parameters (mainly myocardium material model parameters) are assumed to be known. We thus input key boundary conditions: the pressure, *p*, and the active stress, *T*_*Ca*_, as an internal variable to obtain a displacement field, **u**, using

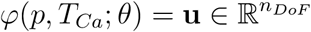

where *θ* represents the network parameters. The PDE is fixed when a hyperelastic material model and fiber field are defined. By fixing the state of the PDE, we are locking in a particular residual operator. In a finite element setting, we want the residual to be zero when the appropriate solution is obtained, so we do the same here by making the residual the loss

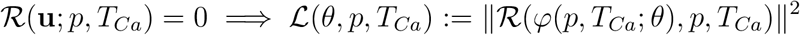

So, we are effectively minimizing the same function used in traditional FE methods, but without the machinery and challenges of the traditional methods, and bring in the ability to train over a wide range of solutions simultaneously and utilize much of the helpful functional features of traditional FE methods.

Training starts with samples taken over a set S representing the physiological range of the parameters, *p* ∈ [0, 120] mmHg and *T*_*Ca*_ ∈ [0, 60] kPa. Thus for each training step, samples are drawn, the network predicts the corresponding solution, and the solution is pushed through the traceable finite element residual calculation for the loss function.

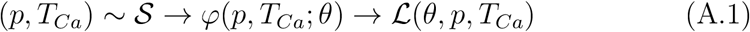

(*p, T*_*Ca*_) ∼ S → *ϕ*(*p, T*_*Ca*_; *θ*) → L(*θ, p, T*_*Ca*_) (A.1) By using the automatic-differentiation capabilities, the residual is differentiated with respect to *θ* to compute the value and gradient of A.1 which is JIT-compiled. This function is then used for optimizing the parameters of the neural network to minimize the residual. This allows training the parameterized PDE over desired values in a single run without requiring data.

## Appendix B Prolate Spheroid Description

To setup the mesh and fibers for the prolate spheroid, we use a mesh generation function that’s contained in `CARDIAX`. The generated mesh gives the appropriate geometry as well as surface tags for applying boundary conditions. Then we use a Laplace-Dirichlet Rule-Based method to obtain the fiber field. The exact fiber field will depend on the specific method used, but we obtain fibers as depicted in ?? that rotate through the ventricle wall.

The PDE being solved is a hyperelastic material model which is the same as the one used in [14]. The strain energy function is split into deviatoric and volumetric componenets

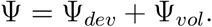

The deviatoric component is given by a transerve isotropic form where fibers are defined

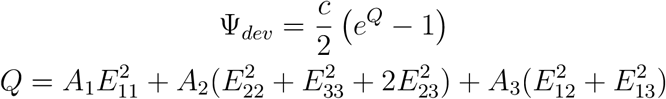

where *E*_*ij*_ = v_*i*_ · Ev_*j*_ is the Green-Lagrange strain contracted in the corresponding fiber, sheet, and normal directions. The volumetric component is given by

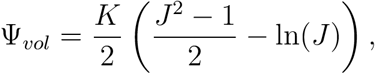

where *J* = det(F) is the determinant of the deformation gradient. The total stress is given by adding the active stress component which is defined by

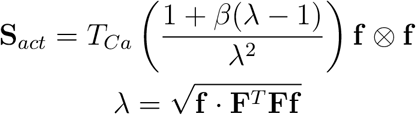

where f is the fiber direction and *T*_*Ca*_ is the magnitude of active stress.

